# Evaluating Cross-Platform Batch Correction Methods for Integrated Microarray and RNA-seq Data Analysis

**DOI:** 10.1101/2024.09.30.615938

**Authors:** Xuejun Sun, Yu Zhang, Chuwen Liu, Xiaojing Zheng, Fei Zou

## Abstract

Integrated analysis of gene expression data across studies is important for understanding complex traits, but combining microarray and RNA-seq data remains challenging because of substantial platform-related technical differences. In this study, we evaluated ten commonly used batch-effect correction methods for cross-platform integration and classified them into three groups representing a spectrum from least to most data processing: unsupervised sample-wise methods, unsupervised gene-wise methods, and supervised methods. Unlike the first two groups, supervised methods use outcome information to guide correction. Performance was assessed for distribution alignment, clustering, outcome prediction, and differential expression (DE) analysis using three paired real microarray–RNA-seq datasets and simulation studies. Supervised methods achieved strong distribution alignment and clustering by biological group, but they introduced information leakage and inflated Type I error, making them unsuitable for unbiased discovery. Among unsupervised methods, sample-wise approaches were often conservative under balanced settings. They showed inflated Type I error under imbalanced settings. In contrast, gene-wise methods generally maintained appropriate Type I error control and achieved higher power, with `limma` showing the strongest DE performance. Meta-analysis that combines p-values also controlled Type I error well and provided competitive power. For outcome prediction, both sample-wise `QN` and gene-wise `limma` performed well, with `limma` providing the strongest overall performance and `QN` offering the practical advantage that new samples can be normalized to an existing reference distribution without refitting. Overall, `limma` is recommended as the best general-purpose method for cross-platform integration.

## Introduction

High-throughput gene-expression profiling has become a fundamental tool for studying transcriptome-wide changes associated with disease, supporting the discovery of key genes, biological pathways, and molecular biomarkers (Hsu et al., 2019; Tomfohr et al., 2005). The rapid expansion of publicly available gene-expression repositories, including GEO, TCGA, and CCLE (Barretina et al., 2012; Barrett et al., 2012; The Cancer Genome Atlas Research Network et al., 2013), has created new opportunities for integrative cross-study analyses, which can improve statistical power and enhance the robustness and reproducibility of molecular signatures. However, substantial batch effects and technical heterogeneity across studies from different platforms remain major challenges for effective data integration.

Microarray and RNA sequencing (RNA-seq) are the two primary platforms for gene expression profiling, yet they differ fundamentally in their data-generating mechanisms. Microarray technology relies on hybridization of RNA to predefined probes and typically yields continuous measurements with a limited dynamic range (Schena et al., 1995). After log transformation, microarray expression data are often approximately normally distributed, making them amenable to Gaussian-based statistical methods. In contrast, RNA-seq uses high-throughput sequencing to quantify transcript abundance, generating count-based data that are commonly modeled using discrete probability distributions, such as the Poisson or Negative Binomial distributions (Wang et al., 2009; Kukurba and Montgomery, 2015). Beyond distinct statistical properties, cross-platform variations are driven by probe-specific hybridization affinities in microarrays, which alter hybridization efficiency and signal intensity (Koltai and Weingarten-Baror, 2008). The resulting variations systematically skew gene expression rankings across the transcriptome.

Batch-effect correction methods are widely used to reduce unwanted technical variation while preserving biological signal in integrative analyses. These methods differ in both their statistical assumptions and implementation strategies. They can be classified as supervised or unsupervised according to whether biological outcome or group information is used during correction, and as subject-wise or gene-wise depending on whether correction is applied separately to each subject or to each gene. Supervised methods may improve the apparent alignment between datasets, but they can introduce information leakage by reusing outcome information in downstream analyses, potentially leading to biased performance estimates. Although numerous batch-correction methods have been proposed, their relative performance in cross-platform settings remains incompletely understood, particularly in the context of downstream analyses such as clustering, outcome prediction, and differential expression analysis. Existing reviews and comparative studies have largely focused on methodological descriptions or descriptive performance metrics, with limited evaluation of how these approaches influence downstream analytical results in practice. Moreover, the impact of supervised batch-effect correction methods on statistical validity, including Type I error control and statistical power, has not been systematically evaluated.

In this study, we conduct a comprehensive, task-oriented evaluation of ten commonly used batch-effect correction strategies for integrating microarray and RNA-seq data. Using simulated data and three paired real-world cross-platform datasets, we assess method performance across four downstream tasks: distributional alignment, clustering, cross-platform outcome prediction, and detection of differentially expressed genes. For differential expression analysis, we compare two strategies: (1) direct testing using pooled or batch-corrected data, and (2) meta-analysis that combines study-specific results. To avoid information leakage and biased performance estimates, supervised approaches were excluded from downstream prediction and differential expression power analyses. Through this downstream-task-driven evaluation, our results provide practical guidance on when batch-effect correction is necessary and how cross-platform gene-expression data should be analyzed to ensure valid and reproducible inference.

## Methodology of Evaluated Batch Correction Methods

In this study, we evaluate ten batch-effect correction methods that are commonly used or adapted for cross-platform integration of microarray and RNA-seq data. To facilitate systematic comparison, we classify these methods along two dimensions: whether biological outcome or group information is used during correction, distinguishing supervised from unsupervised methods, and whether correction is performed at the subject level or the gene level. Table 1 summarizes the core methodology of each approach.

**Table 1.** Summary of evaluated batch-effect correction methods.

| Method Group | Method | Methodology |
| --- | --- | --- |
| Unsupervised Subject-wise | **Quantile Normalization (QN)** (Bolstad et al., 2003) | **QN** aligns each sample to a common reference distribution. For each sample, expression values are sorted, replaced by the values at the same ranks in the reference distribution, and then returned to the original gene order. |
|  | **Angel's Method** (Angel et al., 2020) | **Angel's method** transforms expression values within each sample into ranks and rescales them to the interval [0, 1], where 1 corresponds to the highest expressed gene and 0 to the lowest. |
| | **Training Distribution Matching (TDM)** (Thompson et al., 2016) | **TDM** transforms RNA-seq data to better match the distribution of microarray data in the $\log_2$-transformed expression space. Using microarray as the training reference, it estimates interquartile-range-based tail ratios, truncates extreme RNA-seq values, and rescales RNA-seq values to the empirical range of the microarray data. |
| Unsupervised Gene-wise | **MatchMixeR (MMR)** (Zhang et al., 2020) | **MMR** estimates platform-specific differences from matched gene expression profiles measured on two platforms. For each gene, it fits a linear mixed-effects model relating one platform to the other, with shared fixed effects and gene-specific random effects, and applies the estimated transformation to convert new data to a chosen reference platform. |
|  | **ComBat** (Johnson et al., 2007) | **ComBat** uses an empirical Bayes framework to estimate and remove batch-specific location and scale effects for each gene while borrowing information across genes to stabilize estimation. It adjusts expression values to a common scale while preserving variation associated with specified biological covariates. |
|  | **RNA Breast Cancer (RNABC)** (Pedersen et al., 2018) | **RNABC** first applies quantile normalization to RNA-seq data using microarray data as the reference distribution. ComBat is then applied to the combined RNA-seq and microarray data to remove batch effects. |
|  | **Shambhala2** (Borisov et al., 2022) | **Shambhala2** converts transcriptomic profiles from different platforms into a fixed universal reference format. It first applies quantile normalization with an auxiliary calibration dataset and then uses cubic normalization (Junet et al., 2021) to map each sample toward a definitive reference dataset, followed by gene-wise rescaling to match the reference mean and standard deviation. |
| | **limma** (Ritchie et al., 2015) | **limma** fits a gene-wise linear model to adjust for batch effects while estimating expression differences. Its empirical Bayes variance moderation shrinks gene-specific residual variances toward a common or trend-dependent variance, improving stability, power, and Type I error control. In this study, RNA-seq data are first $\log_2$ transformed and quantile normalized within platform before correction. |
| Supervised | **Rank-In** (Tang et al., 2021) | **Rank-In** converts within-sample expression values into weighted internal gene-expression ranks, preserving both relative ordering and local expression-intensity trends. It then applies singular value decomposition to estimate latent technical factors and subtracts them to obtain an adjusted matrix. The method uses class labels in downstream differential expression and clustering evaluations. |
|  | **COCONUT** (Sweeney et al., 2016) | **COCONUT** estimates batch effects using control samples with ComBat and then applies the derived correction parameters to case samples. By using known group labels during correction, it can improve alignment in some settings but may introduce information leakage in downstream analyses. |

**Table 2.** Cross-platform prediction performance (AUC) on the METSIM dataset using Lasso regression. Models were trained on one platform and tested on the other after batch-effect correction. The highest AUC in each direction is shown in bold.

| Method | Train array → Test seq | Train seq → Test array |
| --- | --- | --- |
| No Integration | 0.785 | 0.672 |
| QN | 0.821 | 0.889 |
| TDM | **0.887** | 0.778 |
| Angel | 0.773 | 0.880 |
| MMR | 0.781 | 0.672 |
| Combat | 0.827 | 0.850 |
| RNABC | 0.812 | 0.786 |
| Shambhala2 | 0.850 | 0.869 |
| limma | 0.808 | **0.898** |

Unsupervised subject-wise methods make relatively modest adjustments and do not fit explicit gene-wise models. Instead, they correct batch effects at the sample level by aligning overall expression distributions across subjects. In contrast, unsupervised gene-wise methods correct each gene separately and often model platform-specific effects directly; some methods further borrow information across genes, for example through empirical Bayes adjustment, to stabilize estimation and improve correction. Supervised methods use the most information because they incorporate biological group or outcome labels during correction. Although this can produce stronger apparent alignment, it may also introduce information leakage and lead to biased downstream results.

These classes of methods are expected to differ in both statistical performance and practical utility. Subject-wise methods are generally simple to implement and can be more convenient when new samples arrive, but they may not adequately remove gene-specific platform effects. Gene-wise methods often provide stronger control of gene-level technical variation and may therefore be better suited for inference tasks such as differential expression analysis. Supervised methods may improve apparent agreement across platforms, especially in clustering, but their use of outcome information can compromise valid downstream evaluation. This motivates a systematic comparison across multiple downstream tasks.

## Method and Data

A unified evaluation framework was used to benchmark ten batch-effect correction methods for integrating microarray and RNA-seq data. The framework assesses how different correction strategies affect downstream analyses commonly performed in cross-platform gene expression studies. Specifically, methods were evaluated based on distributional alignment, sample clustering, cross-platform outcome prediction, and differential expression (DE) detection in simulated data and real data.

### Simulation data analysis

Simulation studies are conducted to evaluate statistical properties that cannot be reliably assessed using real data alone, including Type I error control, statistical power, and outcome prediction. Simulated datasets are generated by resampling paired microarray and RNA-seq profiles with replacement from control samples in the METSIM (Subsection 3.3.3) study, thereby preserving realistic cross-platform dependence while minimizing biological heterogeneity.

For differential expression (DE) evaluation, two analysis strategies are considered. In the joint-analysis strategy, corrected microarray and RNA-seq samples are pooled and DE p-values are calculated using the Wilcoxon rank-sum test on the combined data. In the meta-analysis strategy, DE p-values are first calculated separately in the microarray and RNA-seq datasets using the Wilcoxon rank-sum test and then combined using the Cauchy method. Comparing these strategies allows us to assess when direct joint analysis is sufficient and when meta-analysis provides better Type I error control or statistical power.

Cauchy-based meta-analysis combines statistical evidence across multiple studies by aggregating p-values while explicitly accommodating dependence among them (Liu and Xie, 2020). Specifically, let *p*_*g*1_, … , *p*_*gK*_ denote the p-values for gene *g* obtained from *K* related studies. The Cauchy combination test constructs the statistic

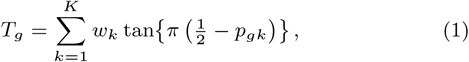

where *w*_*k*_ ≥ 0 are prespecified weights satisfying 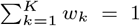. Under the global null hypothesis that gene *g* is not differentially expressed in any study, *T*_*g*_ follows an approximate standard Cauchy distribution regardless of the dependence structure among the *p*_*gk*_. The combined p-value for gene *g* is computed as

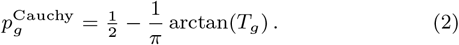

This formulation enables valid meta-analysis without explicitly modeling the correlation among studies.

Under the null setting, no true DE is present. Case–control labels are randomly assigned under four study designs: *Balanced*, in which both microarray and RNA-seq have balanced case–control groups; *Matched Imbalance*, in which both platforms share the same imbalanced case–control ratio (e.g., 3:7); *Reversed Imbalance*, in which the two platforms have opposite imbalance patterns (e.g., 3:7 in microarray and 7:3 in RNA-seq); and *Mixed Balance–Imbalance*, in which one platform is balanced while the other is imbalanced (eg., balanced in microarray and 3:7 case control ratio in RNA-seq). These scenarios are used to evaluate Type I error control and to examine whether batch-effect correction induces spurious sample clustering, particularly for supervised methods that may introduce artifacts through outcome-guided adjustment.

To illustrate the behavior of supervised batch-effect correction under the null setting, we applied Rank-in to simulated data with no truly DE genes. Case and control labels were randomly assigned under each balance design, and PCA plots were generated before and after correction.

Under the alternative setting, artificial DE signals are introduced by designating 2,000 DE genes, including 1,000 upregulated and 1,000 downregulated genes, among 16,160 total genes. Statistical power is then evaluated across varying sample sizes under the *Balanced* design, varying effect sizes under the *Balanced* design, and all four balance designs while holding sample size and effect size fixed.

Cross-platform outcome prediction performance is evaluated using Lasso regression implemented in the `glmnet` package in R. Models are trained on one platform and tested on the other. The regularization parameter is selected by 5 fold cross-validation on the training platform, after which the final model is fit using all training samples and applied to the other platform for prediction. This reflects practical cross-platform applications, such as influenza vaccine studies in which a predictor developed from historical microarray cohorts is applied to RNA-seq datasets. Performance is evaluated under the *Balanced* and *Matched Imbalance* designs using the area under the Receiver Operating Characteristic curve (AUC). The *Reversed Imbalance* and *Mixed Balance–Imbalance* settings are not considered for cross-platform prediction because differing case–control ratios across platforms would introduce confounding and make prediction performance difficult to interpret.

### Real data analysis

Real-data analyses were conducted using paired cross-platform datasets to assess whether batch-effect correction methods reduce platform-specific effects while preserving biologically meaningful signals. Because the microarray and RNA-seq profiles were measured from the same underlying samples, these paired data minimize biological differences while retaining platform-related variation. For each dataset and correction method, batch correction was first applied to the paired microarray and RNA-seq data. All corrected gene-expression values were then pooled across samples within each platform, and histograms were drawn to compare the overall microarray and RNA-seq expression distributions. This qualitative assessment evaluates whether each correction method improves cross-platform distributional alignment while avoiding excessive distortion of the expression profiles.

Next, the sample-level structure is evaluated using hierarchical clustering based on Euclidean distance. Clustering patterns are examined with respect to both biological outcome groups (e.g., case versus control) and platforms (e.g., microarray versus RNA-seq). An effective correction method is expected to reduce clustering driven by the platform while preserving clustering by biological outcome.

Finally, cross-platform outcome prediction is evaluated using Lasso regression under the same framework as in the simulation study, using the METSIM (Subsection 3.3.3) data. Prediction models are trained on harmonized microarray data and tested on harmonized RNA-seq data, and vice versa. Model tuning is performed using 5-fold cross-validation on the training platform; after the regularization parameter is selected, the final model is fit using all training samples and applied to the other platform for prediction. Predictive performance is quantified by the AUC, given the matched class imbalance in the real dataset.

### Real datasets were used in this study

#### MAQC/SEQC data

This dataset was obtained from the SEQC/MAQC project (GEO accession numbers: GSE56457 and GSE47774) (SEQC/MAQC-III Consortium, 2014; Su et al., 2014), a large-scale initiative led by the U.S. Food and Drug Administration (FDA) to evaluate the reliability and reproducibility of microarray and next-generation sequencing technologies. We focused on two well-characterized reference samples: Sample A, a pooled cell-line mixture, and Sample B, a human brain reference sample, which were treated as the case and control groups. Both samples were profiled on microarray and RNA-seq platforms. To support a balanced cross-platform comparison, we included equal numbers of profiles for each sample type, specifically 16 RNA-seq profiles and 8 microarray profiles for both Sample A and Sample B.

#### CCLE data

This dataset contains 107 microarray and 105 RNA-seq expression profiles from cancer cell lines in the Cancer Cell Line Encyclopedia (CCLE) (The Cancer Genome Atlas Research Network et al., 2013). Profiles from the two platforms are treated as paired because they were generated from the same underlying cell lines and therefore reflect the same biological state measured by different technologies. From this dataset, we selected large intestine and breast cancer cell lines, which were treated as the case and control groups, respectively.

#### METSIM data

The METSIM study is a cohort investigation focused on the relationship between adipose gene expression and cardio-metabolic traits (El-Sayed Moustafa et al., 2020; Civelek et al., 2017). Both microarray and RNA-seq data from this study are publicly available under the GEO accession numbers GSE135134 and GSE70353, derived from the same subcutaneous adipose tissue samples. For our analysis, we select 660 paired microarray and RNA-seq samples. To define case and control groups, we use the Matsuda Index, a measure of insulin sensitivity. Participants with a Matsuda Index of ≥ 4 are classified as controls, while those with a Matsuda Index of *<* 4 are classified as cases.

## Results

### Simulated data results

To assess Type I error control under the null hypothesis, we performed no-DE-gene simulation studies across four balance designs and both small and large sample sizes for all 10 batch-correction methods and the meta-analysis approach. As shown in Figure 1, the unsupervised subject-wise methods (`QN`, `Angel`, and `TDM`), together with `MMR`, were generally conservative under the *Balanced* and *Matched Imbalance* designs at both sample sizes, but exhibited inflated Type I error under the *Reversed Imbalance* design and under the large-sample *Mixed Balance–Imbalance* design. Among the unsupervised gene-wise methods, `ComBat`, `RNABC`, and `limma` generally maintained Type I error close to the nominal 0.05 level, whereas `Shambhala2` showed mild inflation in some large-sample *Reversed Imbalance* settings and `ComBat` showed inflation in the small-sample *Reversed Imbalance* setting. The supervised methods were less robust: `Rank-In` showed substantial Type I error inflation across all conditions, while `COCONUT` was mainly inflated under the *Reversed Imbalance* design. In contrast, the meta-analysis approach consistently controlled Type I error well across all scenarios.

**Figure 1.**
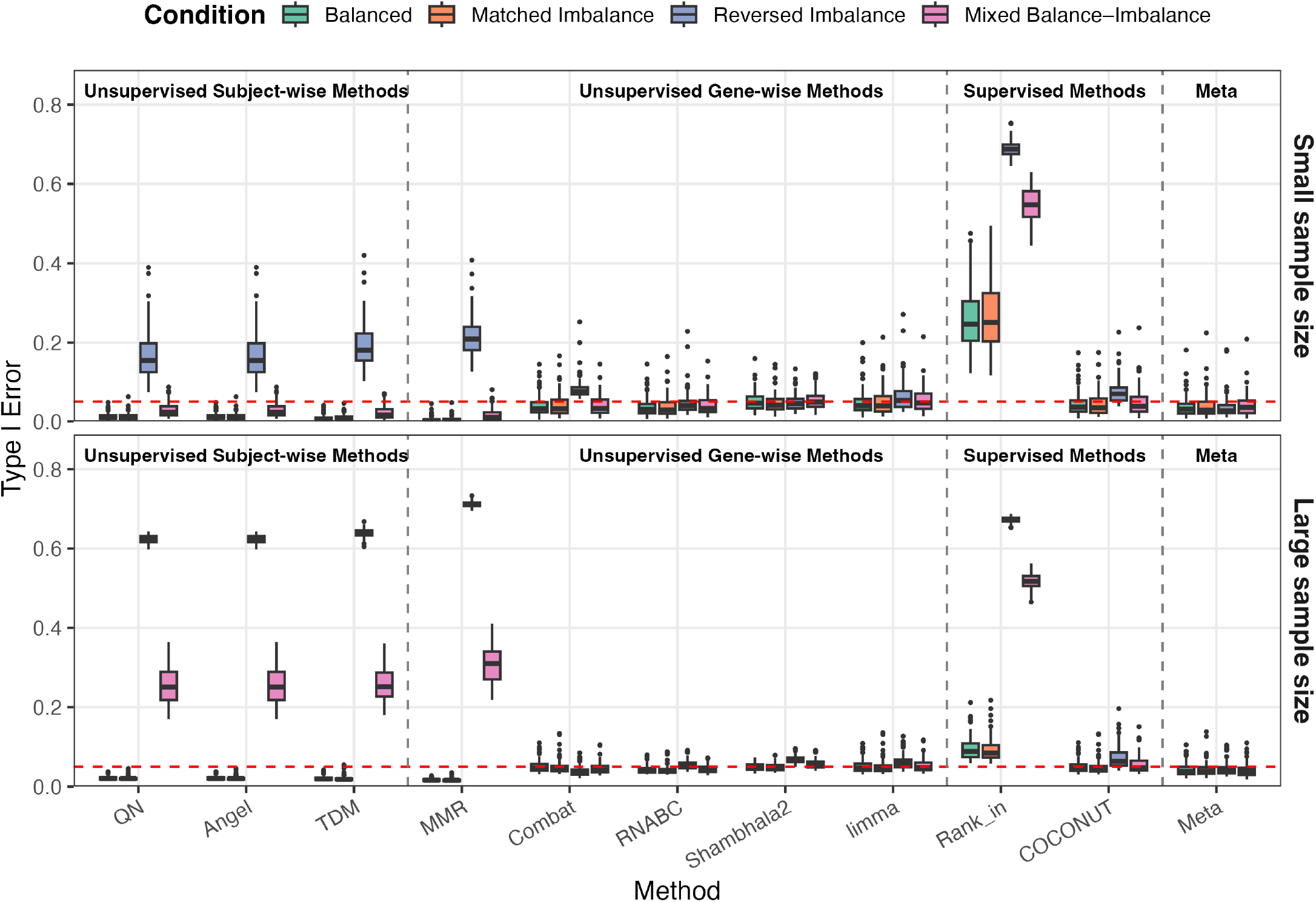
Type I error under the null setting across batch-effect correction methods for a small-sample setting (*n* = 20 for microarray and *n* = 20 for RNA-seq) and a large-sample setting (*n* = 100 for microarray and *n* = 100 for RNA-seq). Results are shown under four label-balance designs: *Balanced, Matched Imbalance, Reversed Imbalance*, and *Mixed Balance–Imbalance*.

As illustrated in Figure 2, using outcome-group information during batch-effect correction can introduce artificial structure under the null setting. Before correction, samples are primarily separated by platform, while the randomly assigned case and control labels do not form meaningful clusters. After applying `Rank-in`, however, the corrected data show apparent separation by case–control group, particularly under the imbalanced designs, even though no truly differentially expressed genes are present. This suggests that Rank-in can transfer the supplied group labels into the corrected expression matrix, creating spurious biological patterns and increasing the risk of false-positive findings in downstream analyses.

**Figure 2.**
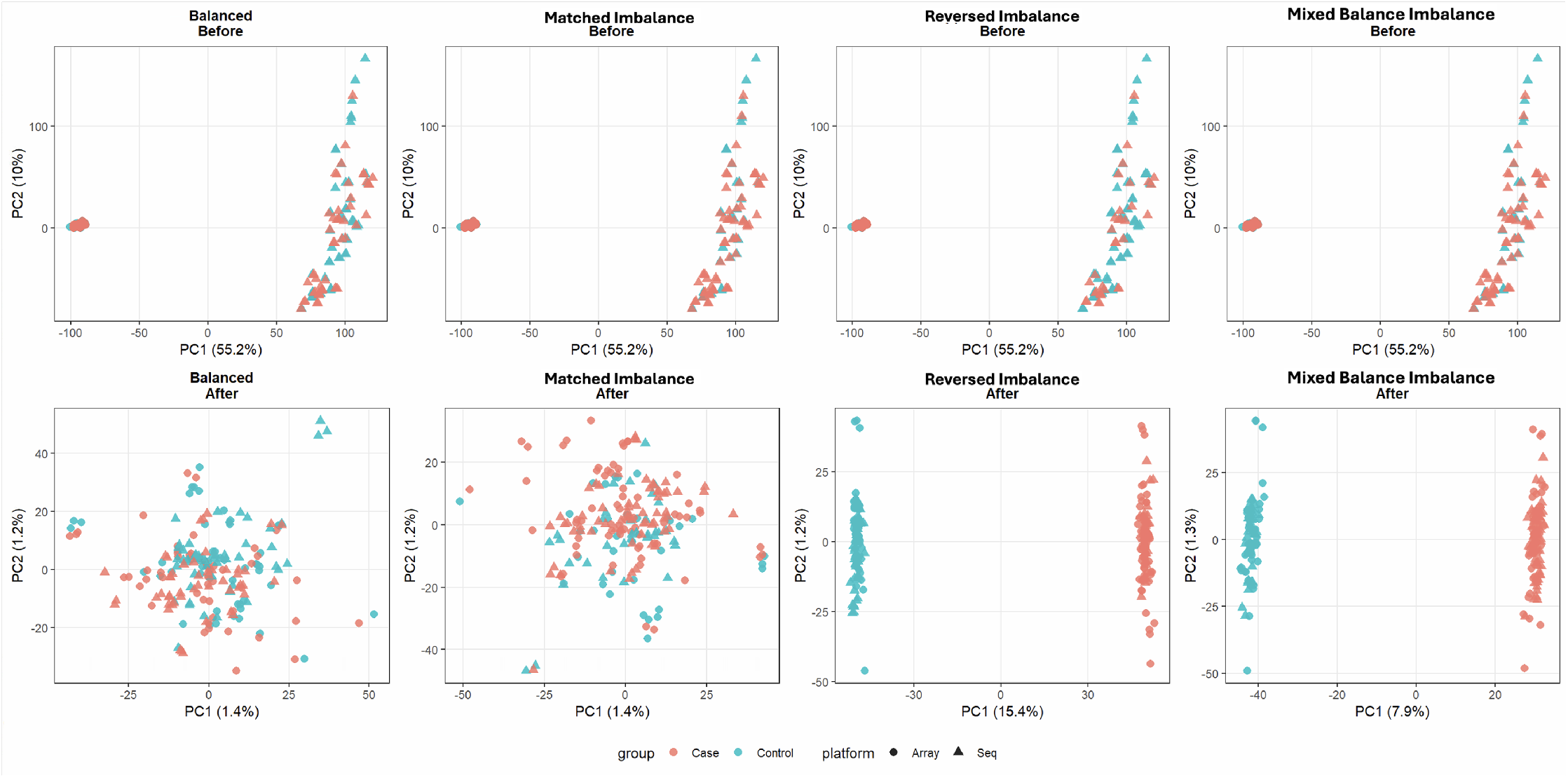
Sample clustering result for Rank-in under the simulation null setting with no truly differentially expressed genes. Case and control labels were randomly assigned, batch correction was performed using Rank-in, and the corrected data were plotted to assess clustering patterns by case–control status and platform.

To evaluate statistical power under the alternative hypothesis, we introduced artificial differential expression signals and assessed power across varying sample sizes, varying effect sizes, and the four balance designs. Supervised methods were excluded from this evaluation because they use outcome information during correction, which can lead to information leakage.

As shown in Figure 3, the sample-size and effect-size analyses were conducted under the *Balanced* design. For the sample-size analysis, the unsupervised subject-wise methods consistently showed lower power than most gene-wise methods. Among the gene-wise methods, `MMR` performed the worst across nearly all settings, whereas `ComBat`, `RNABC`, `Shambhala2`, and `limma` achieved substantially higher power. `Limma` showed the highest power across all sample sizes. `Shambhala2` and the meta-analysis approach were comparable to `limma` at larger sample sizes, although both showed lower power in small-sample settings.

**Figure 3.**
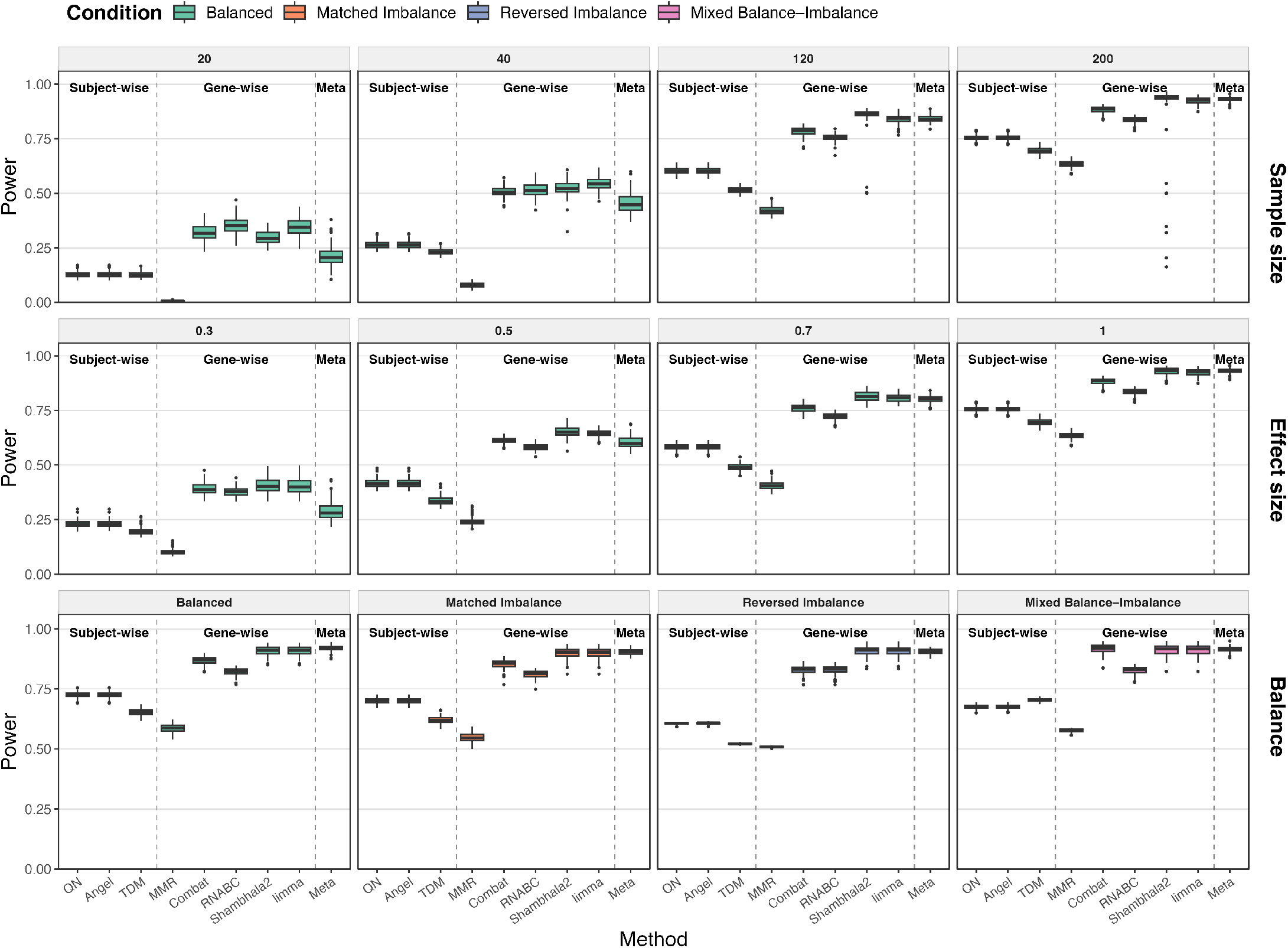
Statistical power under the alternative setting across batch-effect correction methods. Power is evaluated by sample size (top row), effect size (middle row), and balance design (bottom row). In the sample-size analysis, total sample sizes of 20, 40, 120, and 200 are considered, where the total denotes the combined number of microarray and RNA-seq samples, under the Balanced design with effect size 1. In the effect-size analysis, effect sizes of 0.3, 0.5, 0.7, and 1 are considered under the Balanced design with total sample size 200. In the balance-design analysis, results are shown under the Balanced, Matched Imbalance, Reversed Imbalance, and Mixed Balance–Imbalance settings with total sample size 200 and effect size 1.

A similar pattern was observed in the effect-size analysis. Subject-wise methods remained less powerful across the range of effect sizes, and `MMR` again showed poor performance. In contrast, `limma` and `Shambhala2` consistently achieved the highest power, while the meta-analysis approach became increasingly competitive as the effect size increased.

For the balance-design analysis, the subject-wise methods and `MMR` were more adversely affected by imbalance and showed reduced power under imbalanced settings. By contrast, the other gene-wise methods were comparatively robust across the different balance designs. Overall, `limma` was the strongest performer across sample size, effect size, and balance scenarios, providing the best combination of high and stable power.

To evaluate prediction performance after batch effect correction, we trained models on one platform and tested them on the other. We focused on gene-wise and subject-wise unsupervised batch correction methods and excluded supervised methods to avoid potential data leakage. Performance was evaluated using AUC under balanced and matched-imbalance conditions with both small and large effect sizes, while keeping the total sample size fixed. Results without any batch correction were also included as the baseline.

Nearly all correction methods improved cross-platform prediction performance compared with the baseline, except for `MMR` Figure 4. Among the subject-wise methods, `QN` achieved the best performance across all scenarios. Among the gene-wise methods, `limma` performed best when training on microarray data and testing on RNA-seq data. When training on RNA-seq data and testing on microarray data, `Shambhala2` performed best, although `limma` showed comparable performance.

**Figure 4.**
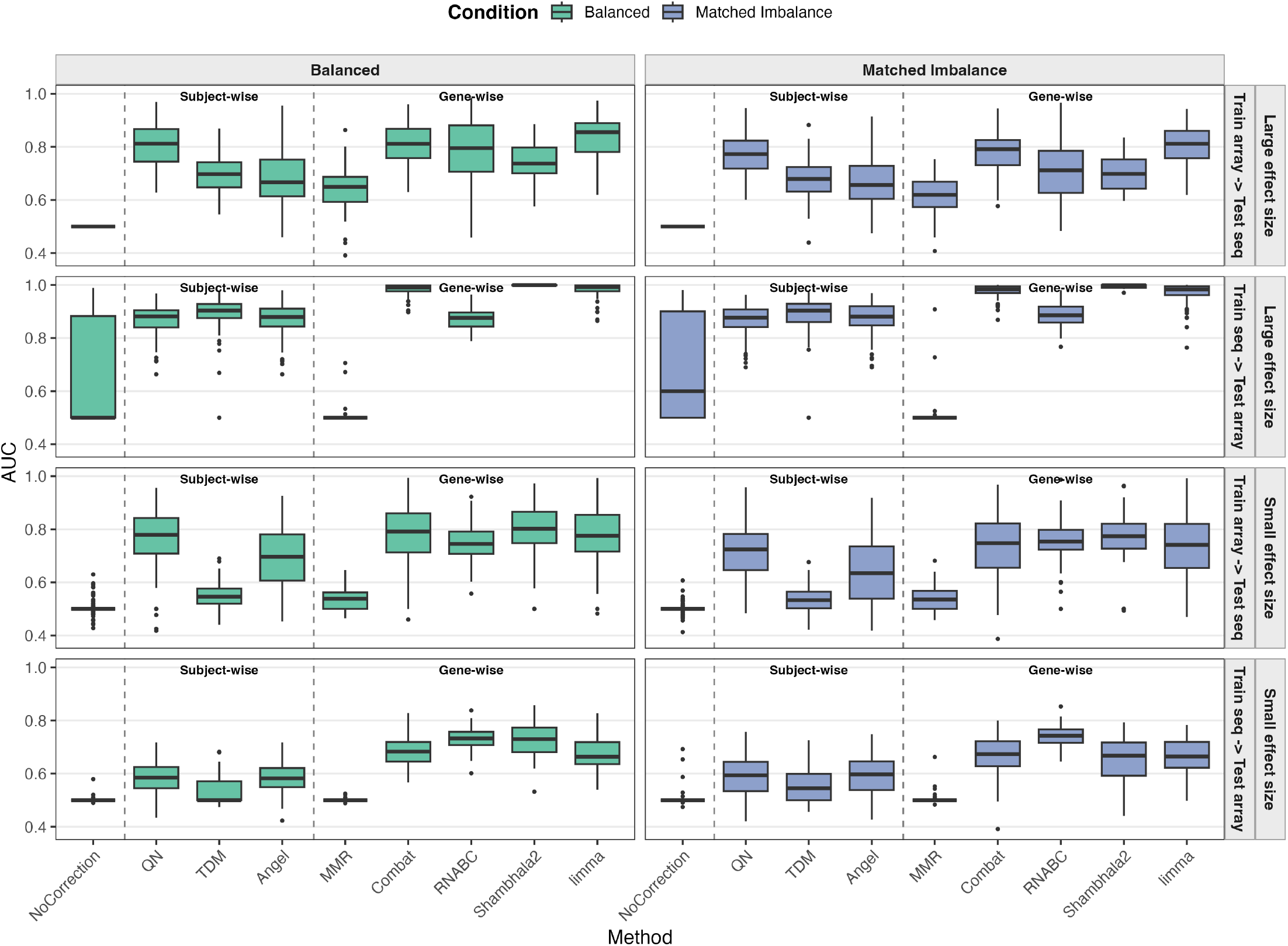
Cross-platform outcome prediction performance measured by AUC under two balance designs (*Balanced* and *Matched Imbalance*), two transfer directions (train on microarray and test on RNA-seq, and train on RNA-seq and test on microarray), and two effect-size settings (*Small effect size* and *Large effect size*). Methods are ordered as no correction, subject-wise normalization methods, and gene-wise normalization methods.

In summary, the unsupervised subject-wise methods were generally conservative under the Balanced and Matched Imbalance settings, which led to lower power for DE detection, and they often failed to adequately control Type I error under more severe imbalance settings. For cross-platform prediction, however, several subject-wise methods remained competitive, with `QN` and `TDM` showing strong performance in some settings. In contrast, the unsupervised gene-wise methods, with the exception of `MMR`, generally maintained good Type I error control, achieved higher power, and also performed well in cross-platform prediction. Among these, `limma` showed the strongest overall performance, while `Shambhala2` was also highly competitive for prediction. Supervised methods were excluded from the power and prediction evaluations because of potential information leakage, and they showed inflated Type I error in DE analysis. The meta-analysis approach also maintained good Type I error control and achieved competitive power when the sample size or effect size was sufficiently large, although it is not directly applicable to cross-platform prediction.

### Real data results

Real-data analyses were performed to evaluate how batch-effect correction methods affect downstream performance in practical cross-platform settings. Three complementary aspects were considered: distributional alignment, sample-level clustering, and cross-platform prediction. Distributional alignment and clustering results are provided in the Supplementary Materials.

Across the SEQC, CCLE, and METSIM datasets, microarray and RNA-seq data showed clear differences in expression range and distributional shape before correction, reflecting substantial cross-platform heterogeneity. Supervised methods, including `Rank-in` and `COCONUT`, produced the strongest visual alignment across platforms, whereas unsupervised gene-wise methods such as `ComBat`, `RNABC`, `Shambhala2`, and `limma` generally reduced platform differences more effectively than subject-wise methods. `MMR` showed the weakest alignment overall (Figure S1). These results suggest that gene-wise unsupervised methods tend to harmonize distributions better than subject-wise unsupervised methods, although supervised methods can produce stronger apparent alignment.

Hierarchical clustering further showed that the effect of batch correction depended on the relative strength of biological and platform signals (Figure S2). In SEQC, most methods preserved clear separation between groups A and B, suggesting strong biological signal except for `Shambhala2`. In CCLE, correction improved separation between breast cancer and large intestine samples for several methods, especially `COCONUT`, `Rank-in`, and `RNABC`, although the stronger clustering from supervised methods should be interpreted cautiously because outcome information is used during correction. In METSIM, separation between control and case samples remained weak before and after correction, suggesting that platform-related variation dominated the biological signal.

Using the METSIM dataset, cross-platform prediction performance was evaluated by AUC when training on one platform and testing on the other. Overall, corrected data outperformed uncorrected data. Among subject-wise methods, `QN` and `TDM` showed strong prediction performance, whereas `MMR` was consistently weakest. Among gene-wise methods, `ComBat`, `Shambhala2`, and `limma` achieved the highest AUC values overall, with `limma` performing best when training on RNA-seq and testing on microarray, and `TDM` performing best when training on microarray and testing on RNA-seq. Overall, gene-wise correction methods tended to provide the most reliable prediction performance, although `QN` and `TDM` were also strong practical options in this real-data analysis.

Overall, the real-data analyses showed that correction performance depended on the downstream task. Supervised methods produced the strongest apparent alignment and, in some datasets, clearer clustering, but these results should be interpreted cautiously because they use outcome information. Among unsupervised methods, gene-wise approaches generally reduced cross-platform differences more effectively and gave the most reliable overall performance, whereas `MMR` was consistently the weakest. For cross-platform prediction, corrected data outperformed uncorrected data, with `ComBat`, `Shambhala2`, and `limma` performing best overall, while `QN`and `TDM` also remained competitive.

## Discussion

Cross-platform integration of bulk transcriptomic data remains challenging because microarray and RNA-seq measurements differ substantially in their measurement scales, data distributions, and sources of technical variation. Paired microarray–RNA-seq data from both simulations and real datasets allowed us to disentangle biological variation from platform-specific effects and evaluate batch-effect correction methods under both controlled and real-world conditions.

Our results show that no single batch-effect correction method is optimal for all downstream tasks. Instead, performance reflects a trade-off across methods ranging from less aggressive to more aggressive correction, namely unsupervised subject-wise methods, unsupervised gene-wise methods, and supervised methods. Subject-wise methods make relatively modest adjustments and were competitive for cross-platform prediction, with `QN` emerging as a practical and effective choice, but they were often conservative under balanced settings and less reliable for DE analysis. Gene-wise unsupervised methods provided the most balanced overall performance, generally maintaining good Type I error control, achieving higher power, and performing well in prediction; among them, `limma` was the most consistently strong performer across simulation and real-data analyses, while `Shambhala2` also showed high power and strong prediction performance, albeit with greater computational cost. Supervised methods, which apply the most aggressive correction by incorporating outcome information, often produced the strongest apparent distributional alignment and clustering, but at the cost of information leakage and inflated Type I error. The meta-analysis approach also performed well, with robust error control and competitive power for DE analysis, although it is not applicable to clustering or cross-platform prediction. Together, these findings suggest that stronger correction does not necessarily yield better overall downstream performance: `limma` is the strongest general-purpose choice, `QN` is a practical option for prediction and incremental data settings, and meta-analysis is preferable for DE analysis.

Beyond statistical accuracy, practical utility also matters. Although `limma` showed the strongest overall performance, it and other gene-wise methods generally need to be refit when new data are added. Subject-wise methods are easier to update because new samples can be normalized against an existing reference distribution, which is advantageous in continuously growing studies; among them, `QN` was especially attractive because it also performed well for prediction. Supervised methods may improve apparent alignment or clustering, but they are less suitable when unbiased inference is required because they use outcome information and may introduce data leakage. Meta-analysis is particularly useful for DE analysis in incremental settings, since each dataset can be analyzed separately and combined later, but it does not extend naturally to clustering or cross-platform prediction.

This study has several limitations. First, the simulations focused on paired integration of two studies from two platforms, whereas real applications may involve multiple studies from the same platform or from multiple platforms, such as microarray, RNA-seq, and other omics technologies, along with additional sources of batch variation and more complex study designs. Second, the real-data prediction analysis was based on one modeling framework, and alternative classifiers or feature-learning strategies may lead to different relative rankings. Third, the current evaluation focused on bulk transcriptomic data and may not fully generalize to other omics settings.

In summary, paired microarray–RNA-seq data provide a useful framework for benchmarking batch-effect correction methods. The results suggest that limma is the strongest overall unsupervised method when robust inference and high power are both required. For DE analysis specifically, meta-analysis is often preferable to direct pooled correction. Therefore, batch-effect correction methods should be selected according to the primary analytical goal rather than assumed to perform uniformly well across all downstream tasks.

## Conflicts of interest

The authors declare that they have no competing interests.

## Funding

This work was supported in part by the National Institutes of Health under grant R01 AI170959-01A1.

## Data availability

The data underlying this article are publicly available from the Cancer Cell Line Encyclopedia (CCLE), the SEQC/MAQC project, and the METSIM study. The SEQC/MAQC data can be accessed through GEO under accession numbers GSE56457 and GSE47774, and the METSIM data can be accessed through GEO under accession numbers GSE135134 and GSE70353. The CCLE microarray and RNA-seq expression profiles are available through the CCLE data portal (https://sites.broadinstitute.org/ccle/) and the corresponding published resource.

## Author contributions statement

X.S., Y.Z., C.L., X.Z., and F.Z. conceived the study. X.S. conducted the experiments, performed the analyses, and drafted the manuscript. Y.Z. and C.L. contributed to data processing, and interpretation of results. X.Z. and F.Z. supervised the study. All authors reviewed, edited, and approved the final manuscript.

## Supplementary Materials

Supplementary Figure S1 shows distributional alignment between microarray and RNA-seq data across the SEQC, CCLE, and METSIM datasets after batch-effect correction. Before correction, the two platforms showed clear differences in expression range and distributional shape, indicating substantial cross-platform heterogeneity. Supervised methods, including Rank-In and COCONUT, generally produced the strongest visual alignment. Among unsupervised methods, gene-wise approaches such as ComBat, RNABC, Shambhala2, and limma reduced platform differences more effectively than subject-wise methods, whereas MMR showed the weakest alignment overall.

Supplementary Figure S2 shows sample-level clustering patterns after correction. In SEQC, most methods preserved separation between groups A and B, suggesting a strong biological signal. In CCLE, several methods improved clustering by biological group, especially COCONUT, Rank-In, and RNABC; however, the stronger clustering from supervised methods should be interpreted cautiously because group information is used during correction. In METSIM, case-control separation remained weak across methods, suggesting that the biological signal was subtle relative to platform-related variation. Overall, these supplementary results show that distributional alignment and clustering are useful diagnostics but should be interpreted together with downstream prediction and differential expression results.

**Figure S1.**
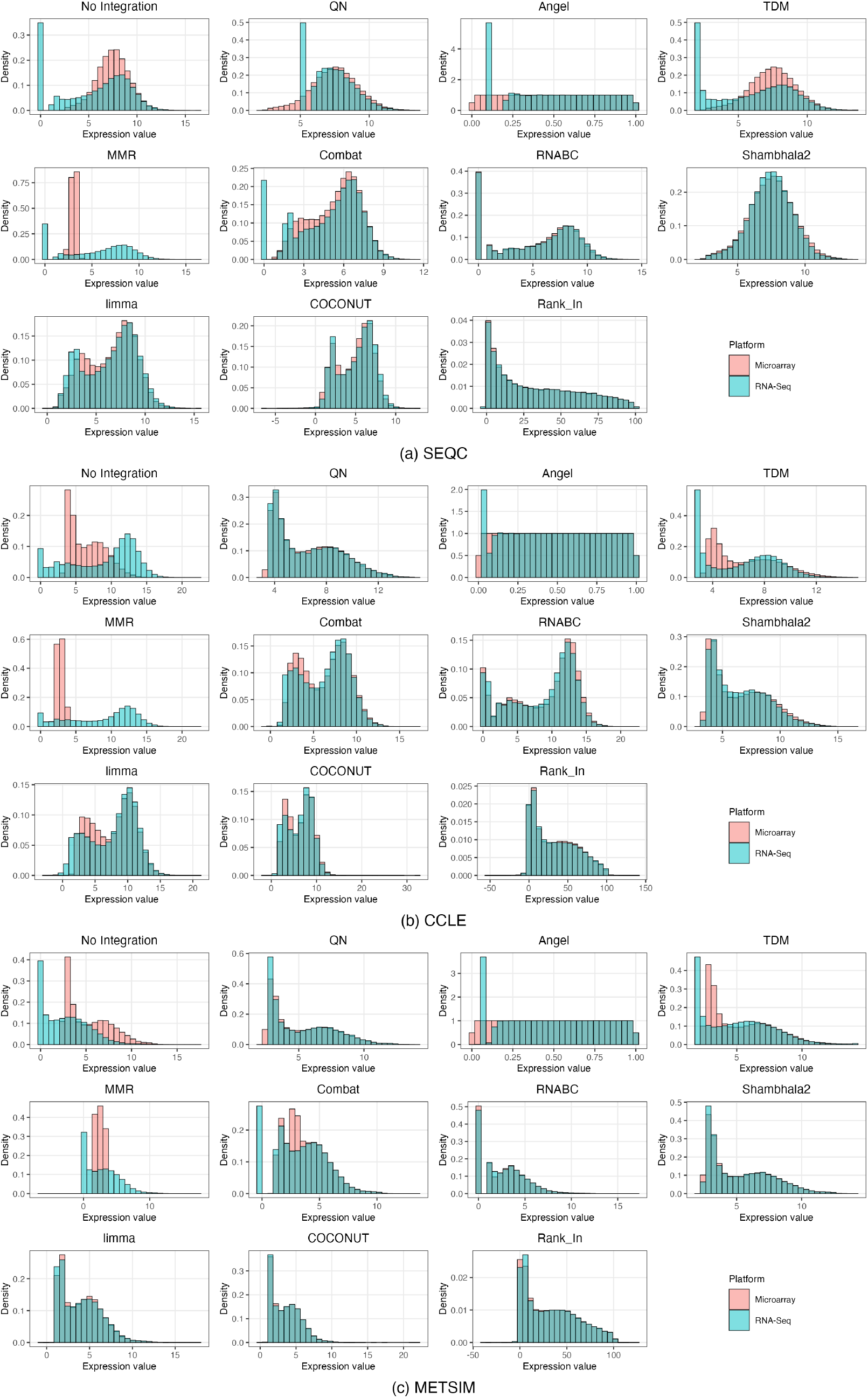
Distribution alignment across three real datasets after batch correction. Histograms are shown for SEQC, CCLE, and METSIM under no integration and multiple correction methods, with microarray and RNA-seq samples overlaid to assess cross-platform harmonization.

**Figure S2.**
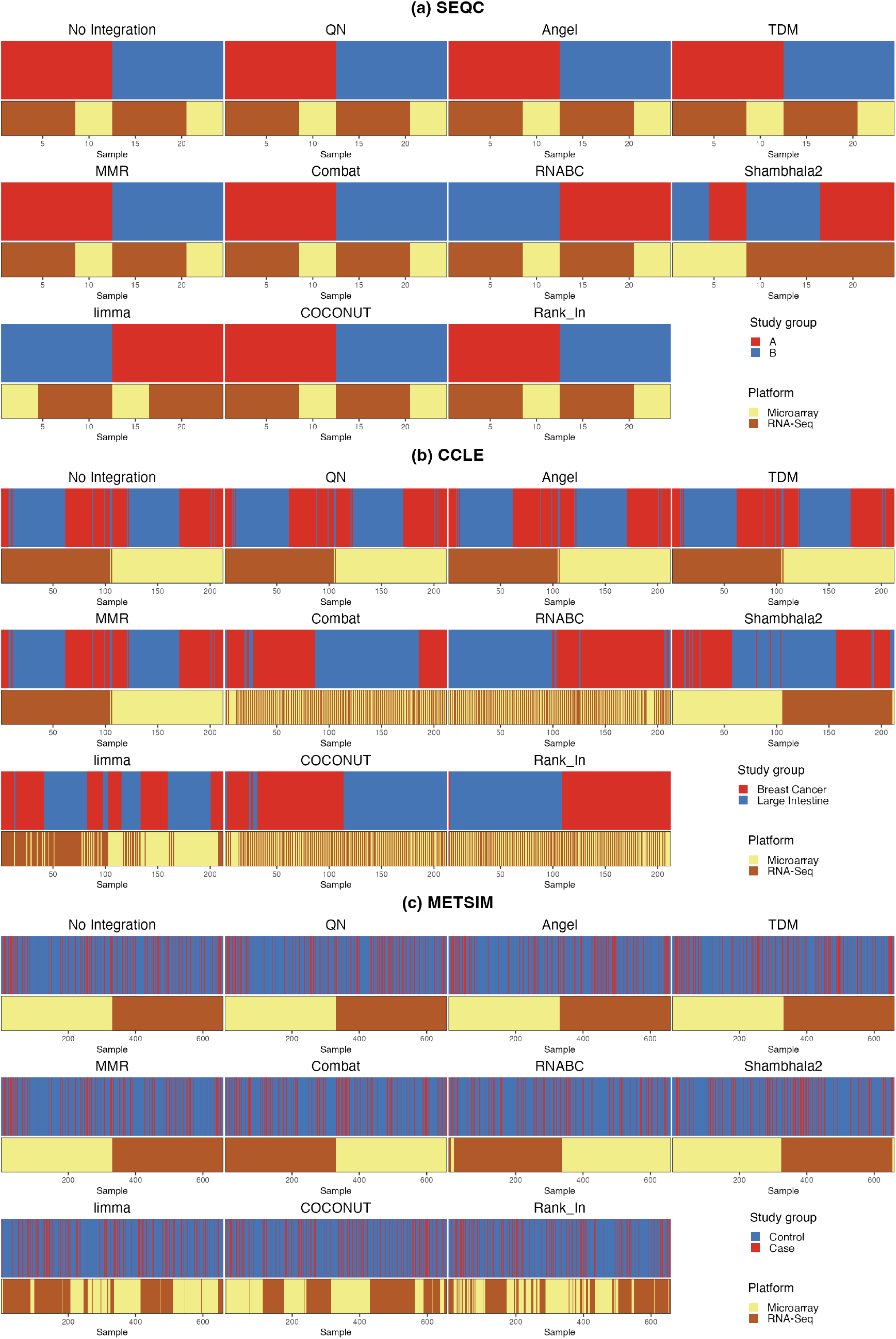
Sample clustering patterns across three real datasets after batch correction. For each method, the top bar indicates biological group labels and the bottom bar indicates platform labels, with samples ordered by hierarchical clustering based on Spearman correlation.

## Notes

### Competing Interest Statement

The authors have declared no competing interest.

### Summary of Updates

Adding one comparing method and manuscript

